# A bioactive soluble recombinant mouse LIGHT promotes effective tumor immune cell infiltration delaying tumor growth

**DOI:** 10.1101/2024.07.30.605649

**Authors:** Maria-Luisa del Rio, Giovanna Roncador, Juan Carlos Cubria, Pascal Schneider, Jose-Ignacio Rodriguez-Barbosa

## Abstract

The TNF family member LIGHT (TNFSF14) binds to two receptors, HVEM (TNFSFR14) and LTβR (TNFSFR3). HVEM functions as a costimulatory molecule, whereas LTβR is involved in the development of lymph nodes and ectopic tertiary lymphoid structures at chronic inflammation sites. The classical approach of fusing soluble recombinant proteins to the Fc fragment of IgG resulted in a functionally inactive Ig.mouse (m) LIGHT protein. However, in line with the fact that TNF family members cluster receptors as trimers, addition of a small homotrimeric domain (foldon) N-terminal of mLIGHT produced an active Ig.Foldon-mLIGHT protein able to bind and engage HVEM and LTβR in a reporter cell-based bioassay.

In the tumor model of B16.F10 melanoma cells implanted into syngeneic recipients, cells transduced with membrane-bound mLIGHT grew as aggressively as mock-transduced cells, but growth of tumors of B16.F10 cells expressing Ig.Foldon-mLIGHT was delayed and characterized by significant immune cell infiltration.

This work unveils the potential of active soluble LIGHT, as a single agent, to recruit cytotoxic cells and dendritic cells at the tumor site to inhibit tumor growth. This effect may be further enhanced with immune checkpoint blockade therapies.

## Introduction

The TNF superfamily comprises 19 type II transmembrane proteins with an N-terminal intracellular region and a conserved trimeric C-terminal extracellular domain known as the TNF homology domain (THD) (1). One of these proteins is LIGHT (homologous to lymphotoxin that exhibits inducible transient expression and competes with herpes simplex virus glycoprotein D for herpes virus entry mediator (HVEM), a receptor expressed by T lymphocytes). The 239 amino acids-long mouse LIGHT (TNFSF14, CD258) is 74% similar to human LIGHT in the THD and regulates innate and adaptive immune responses by engaging two receptors, the herpes virus entry mediator (HVEM, TNFSFR14, CD270) and the lymphotoxin beta receptor (LTβR, TNFSFR3). LIGHT can be either membrane-bound or released as a soluble cytokine by a metalloprotease (2), (3), (4).

LIGHT is structurally related to lymphotoxins (LTα3 and LTα1β2) and is transiently expressed on the cell surface of activated T cells, activated natural killer (NK) cells, neutrophils, dendritic cells (DC), and other immune innate cells, but not on naive T cells, regulatory T cells, or B cells (5), (6), (7). HVEM is expressed predominantly in hematopoietic cells, including naive CD4 T cells, CD8 T cells, B cells, myeloid cells, endothelial cells and in the epithelium of the mucosa at barrier sites (5), (8), (9). LTβR expression is restricted to endothelial, stromal and myeloid cells, but is not detected on lymphoid cells (10), (11). LTα1β2, which also exists as membrane-bound and soluble molecules, is expressed on activated T cells, B cells, and NK cells and its binding to LTβR promotes lymph node development during embryogenesis. In adults, LTβR contributes to the transition of flat endothelium to high endothelial venules (HEV) in pathological scenarios of chronic inflammation. HEVs are critical structures for the immune cells to gain access to the tumor microenvironment and for the subsequent formation of tertiary lymphoid structures (12), (13).

The engagement of HVEM by LIGHT in *trans* unleashes CD28-independent costimulatory and survival signals on T cells and NK cells present in the surrounding proinflammatory environment and sustain their differentiation towards effector cells (14), (15), (16), (17), (18), (19), (20), (21). LIGHT and LTα1β2 also bind to LTβR on stromal cells to induce the release of chemokines attracting lymphoid and myeloid cells. LTβR signaling increases adhesion molecules on endothelial cells, bolstering leukocyte attachment and subsequent transmigration that is of particular importance for the recruitment of primed anti-tumor CD8 T cells stimulated in the tumor draining lymph nodes (22), (4).

Unfortunately, the classical approach of fusing recombinant proteins to the Fc fragments of IgG resulted in inactive Ig.mLIGHT proteins (2), (23), hampering studies of potential effects of exogenous LIGHT on the immune response. Here, we describe a method of inserting a small homotrimeric domain between the Fc and LIGHT proteins to obtain functionally active Ig.Foldon-mLIGHT that binds HVEM and LTβR. In-situ delivery of Ig.Foldon-mLIGHT by B16.F10 melanoma cells, unlike membrane-bound LIGHT, robustly promoted immune cell infiltration and subsequent control of tumor growth.

## Material and methods

### Plasmids for the expression of genes of interest into eukaryotic cells

**Table 1** contains a list of plasmids used for the expression of the genetic constructs of interest. The extracellular regions of mouse HVEM or mouse LTβR were fused to the transmembrane and intracellular domains of human Fas and cloned into retroviral pMSCV-pgk-Puro vector. Genetic constructs of mouse LIGHT were prepared for protein expression in HEK-293T cells or for transduction in B16F10 melanoma cells. Plasmids were produced in TOP10 or Stbl3 strains of *E. coli*. Plasmids were sequenced by Sanger sequencing.

### Production and purification of recombinant proteins

HEK293T cells were cultured and expanded in adherent 15 cm diameter dishes in DMEM 10% FCS, plus additives (glutamine 2 mM, pyruvate 1mM, HEPES 10 mM, 1X non-essential amino acids, 0.05 mM 2-βME, gentamycin 50 μg/ml). The day before transfection, HEK293T cells were trypsinized and seeded at 12×10^6^ per 15 cm diameter Petri dishes in 20 ml of complete medium. The day after, medium was aspirated and 30 µg of plasmid in 1.5 ml of Opti-MEM was mixed with 90 μl lipofectamine 2000 (Invitrogen, Cat. #11668-019) (i.e. 3 µl of lipofectamine per µg of plasmid), left for 20 minutes, then added to cells. Then, an additional amount of 5.5 ml of Opti-MEM was added to cover the plates. Six hours post-transfection, plates were filled up with 15 ml of Opti-MEM. Five days after transfection, the supernatant was collected, centrifuged, passed through a 0.22 μm filter and purified by affinity chromatography on protein G Sepharose (Biovision) (Ig.mLIGHT, Ig.Foldon-mLIGHT, mBTLA.Ig), whereas Flag-human LIGHT was purified by immunoaffinity chromatography using anti-Flag M2 antibody bound to agarose (Sigma), dialyzed against endotoxin free PBS, filtered through 0.22-μm pore size, and stored at −80°C, until use (Supplementary Figure 1A). Purified recombinant proteins were size-fractionated on 8% acrylamide SDS-PAGE under reducing conditions (1 mM dithiothreitol) and boiled. Gels with five micrograms protein per lane were stained with Coomassie Blue.

### Transduction of B16.F10 melanoma cells and Fas-deficient Jurkat cells, and selection of clones

mHVEM:Fas clone 4 reporter cells were generated in Fas-deficient Jurkat JOM2 clone 6 cells as described (24). For the production of retroviral particles, HEK293T cells were transfected in 10 cm diameter poly-L-lysine precoated adherent Petri dishes. 10 μg of Hit60 MoMuLV (Gag/pol), 1.5 μg of pantropic envelope (pCG-VSV-G) and 10 μg of pMSCV-pgk-Puro plasmid (encoding membrane-bound LIGHT or Ig.Foldon-mLIGHT plasmids) in 450 µl of OptiMEM that were mixed with 64.5 µl of lipofectamine 2000 (Thermofisher) (*i.e.* 3 µl of lipofectamine per µg of DNA), incubated for 20 min and added to cells (Table I). Retroviral-containing supernatants were harvested 48 hours later, filtered at 0.45 μm and fresh polybrene (Sigma H9268) was added to a final concentration of 8 μg/ml and mixed. Serial dilutions of supernatants plated in 24-well plates were added to adherent B16.F10 melanoma cells or to Fas-deficient human Jurkat (JOM2 clone 6) cell pellets and then spun down for 5 minutes at 1.250 rpm and seeded in 24-well plates. Transduced cells were cloned by limiting dilution in the presence of 1 μg/ml of puromycin (AG Scientific) (25). Isolated clones were selected for stable expression of the gene of interest on the cell membrane of the transduced cells (either mHVEM-hFAS or mLTβR-hFAS) by flow cytometry using a rat anti-mouse HVEM antibody (clone 6C9) or a rat anti-mouse LTβR antibody (clone 4H8) (Supplementary Figure 1).

**Table I:**
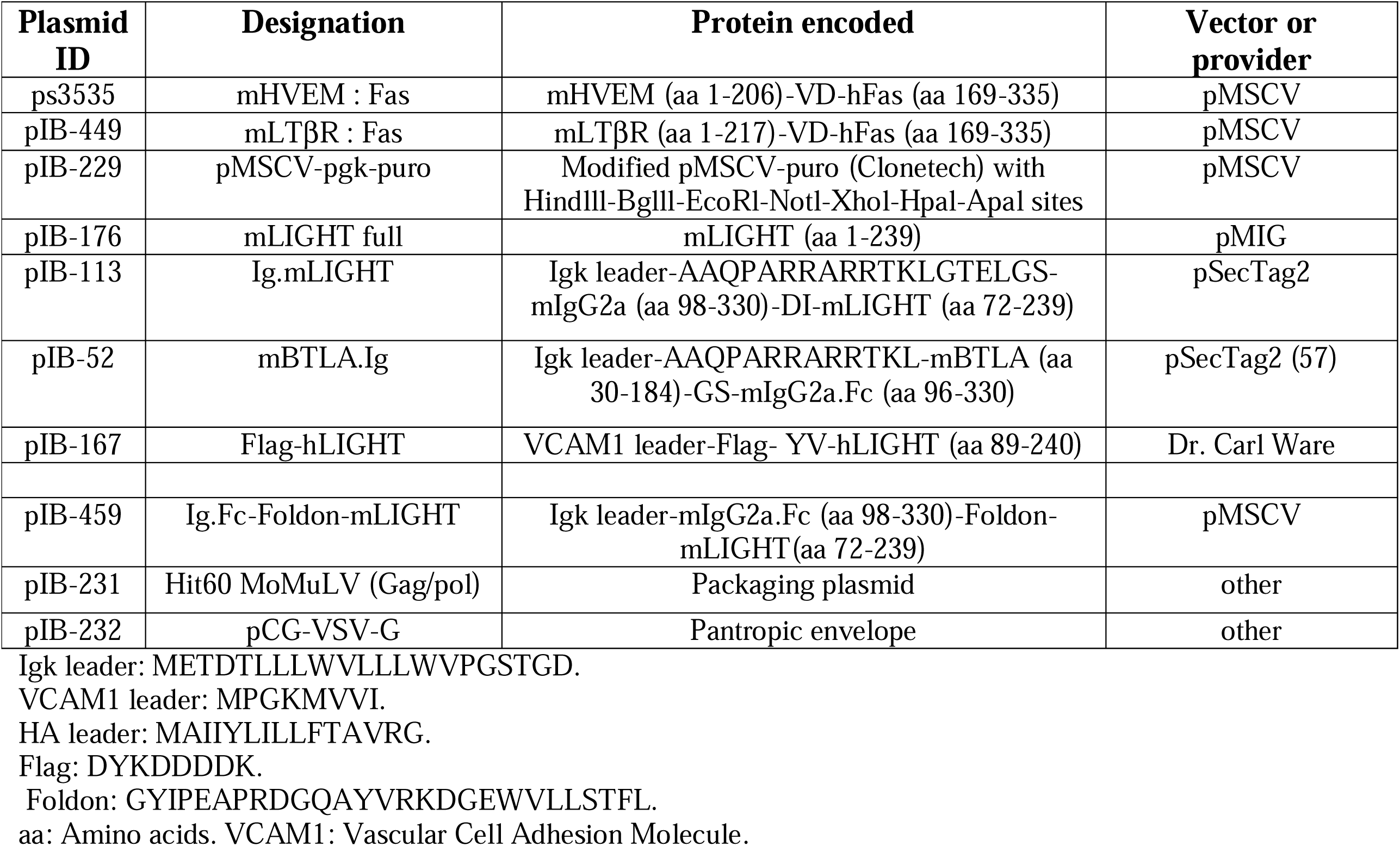
List of plasmids used for the expression of the genetic constructs of interest.

Non-transduced (mock) B16.F10 cells, membrane bound LIGHT- or Ig.Foldon-LIGHT-transduced B16.F10 melanoma cells used in the *in vivo* experiments were cloned by limiting dilution and the resulting single colonies were screened by flow cytometry with anti-LIGHT antibody (clone 3D11) to detect cell-bound LIGHT expression, whereas a sandwich ELISA was implemented for the selection of a cell line secreting recombinant Ig.Foldon-mLIGHT. ELISA plates were coated with a goat anti-mouse IgG2a/b polyclonal antibody (2.5 μg/ml, Nordic) to capture soluble recombinant proteins present in the supernatant of the transduced cells and a biotinylated rat anti-mouse IgG2a (clone R19-15, 1 μg/ml, Beckton Dickinson) was used as detector antibody, following the addition of SA-HRPO (1/10.000). The OD values obtained at 450 nm were extrapolated to the standard curve of known concentrations of mouse IgG2a for the quantification of the recombinant protein in the supernatant.

### Cell viability assay

mHVEM:hFas reporter cells (40.000 cells per well in a final volume of 100 µl) were exposed to titrated amounts of Ig.mLIGHT, Ig.Foldon-mLIGHT, Flag-hLIGHT or mBTLA.Ig for 16 h at 37°C, 5% CO_2._ At the end of this period of incubation, 20 µl of a mixture of 0.9 mg/ml phenazine methosulfate (MTS, Sigma) and 2 mg/ml 3-[4,5-dimethylthiazol-2-yl]-5-[3-carboxymethoxyphenyl]2-[4-sulfophenyl]-2H-tetrazolium (PMS, Promega) (1/20, v/v) was added for a further 2 to 8 h. Absorbance was measured at 490 nm.

The cytotoxic effects of serial dilutions of mock B16.F10 cells or membrane mLIGHT expressing B16.F10 effector cells (starting at 20.000 cells/well) were monitored after 24 h coculture with mHVEM:hFas target cells (40.000 cells/well). Non-adherent mHVEM:hFas target cells were collected and analyzed for cell viability by flow cytometry (FSC/SSC) with a gate for cells that remained alive. In addition, supernatants from mock B16.F10 cells or Ig.Foldon-mLIGHT expressing B16.F10 effector cells (1×10^6^ cells/ml grown during 24 h) were also tested for their cytotoxic activity over mHVEM:hFas target cells (40.000 cells/well), and cell viability was measured at 490 nm using the MTS/PMS assay.

### Mice and tumor implantation and monitoring of tumor growth

Eight- to twelve-week-old C57BL/6J female mice (H-2^b^) were bred in our own animal facility from breeding pairs purchased from Janvier laboratories (France). All animal protocols and experimental designs were authorized and approved by the Animal Welfare Committee of the University of León and endorsed by Castilla and Leon Regional Government (reference # OEBA-ULE-003-2023). Experimental protocols were performed in accordance with the European Guidelines for Animal Care and Use of Laboratory Animals.

B16.F10 melanoma cells at passage 8 were a gift of Dr. Tobias Bald (Bonn, Germany). B16.F10 melanoma cell line is a derivative subline of original B16 melanoma that was selected for its preferential metastatic tumor tropism for the lung when injected intravenously (26). 2 x 10^5^ B16.F10 melanoma cells in 100 µl of PBS were implanted subcutaneously following the technique described by Allard et al., (27). Most tumors exhibit an ellipsoid shape. Tumor volume was monitored every two days from day 8 to day 16 using a digital caliper and tumor volume was calculated with the formula: V = (L×W ^2^) / 2, where length was the greatest longitudinal diameter and width was the greatest transversal diameter.

### Isolation of tumor infiltrating cells

For the isolation of the immune cell infiltrates, tumors were dissected carefully to prevent cross-contamination with brachial and axillar lymph nodes. Dissected tumors were excised finely and digested in 5 ml of digestion buffer HBSS Ca^++^/Mg^++^ without phenol red (containing 5 mg of collagenase D / gr of tumor, 0.5 mg of DNAse I / gr of tumor and supplemented with 25 mM of HEPES pH 7.3, 5 mM CaCl_2_, 5 mM MgCl_2_ and 2% of complement free FCS) and incubated at 37°C for 20 min under shaking conditions at 200 rpm.

Digested tissue was then passed through a nylon mesh holder of 70 microns to a 50 ml tube and spun down. After a washing step with 7 ml of complete RPMI 1640 medium, the pellet was resuspended into 3 ml of Percoll 40% and layered on top of 3 ml of Percoll 80%, and the interphase was collected after centrifugation at 2000 rpm for 20 minutes at RT without break. The isolated cells were washed, resuspended in complete RPMI 1640 medium and spun down. Red blood cells were lysed for 3 min in ACK lysis buffer, cells were washed again, and the pellet was resuspended in 1 ml FACS buffer. One hundred microliters of the cell suspension were incubated with 2 µg/ml of FcR blocker (clone 2.4G2) and then stained with FITC-labeled anti-CD45.2 antibody and the total number of tumor-infiltrating cells were counted in the flow cytometer. The cell counts for that volume were then extrapolated to the final volume of 1 ml in which tumor cells were initially resuspended.

### Flow cytometry

Jurkat cells were collected and resuspended in FACS buffer (PBS containing 2% FCS and 0.5 mM EDTA). Purified Ig.mLIGHT, Ig.Foldon-mLIGHT, Flag-hLIGHT, or mBTLA.Ig fusion proteins (1 μg/ml) were added to the mHVEM:hFas- or LTβR:hFas-deficient Jurkat cells and incubated at 37°C for one hour. Cells were then washed in FACS buffer and incubated with biotinylated anti-mIgG2a (R19-15) or biotinylated anti-Flag (M2) mAbs for 30 min at room temperature. After a washing step, the reactions were developed using SA-Alexa Fluor 647 and the samples were acquired by flow cytometry and collected data analyzed with FlowJo V10 software.

The phenotypic analysis of the tumor-infiltrating leukocytes was performed by flow cytometry using the set of monoclonal antibodies conjugated to fluorochromes listed in **Table 2**. Tumor infiltrating immune cells were first gated on CD45.2-positive cells (pan leukocyte marker) and from there using successive sub-gating strategies, the different immune cell populations were phenotypically characterized. The binding of biotinylated antibodies to cell surface was revealed with streptavidin (SA)-PE, SA-PE/Cy7, SA-Alexa 647 or SA-BV421, depending on the combination of fluorochromes used in each staining. Dead cells and debris were systematically excluded from the acquisition gate by adding propidium iodide (PI). Live cells were gated as PI negative and cellular aggregates were excluded according to FSC-H /FSC-A dot plot profile. Flow cytometry acquisition was conducted on a Cytek® Aurora Spectral Cytometer and data analysis was performed using FlowJo software version 10.

**Table 2:** List of antibodies: specificity, biotin/fluorochrome labeling, clone name and the provider.

| Target | Labelling | Clone | Company |
| --- | --- | --- | --- |
| **Lineage restricted cell surface markers** |  |  |  |
| Isotype rat IgG<sub>2a</sub> | Biotin | AFRC MAC 157 | Home-made (ECACC) |
| Isotype mouse IgG2b | Biotin | MPC-11 | Home made (ECACC) |
| FcγR blocker | unlabeled | 2.4G2 | Home-made (58) |
| CD45.2 | FITC | 104 | Biolegend (#109806) |
| CD11b | PerCP-Cy5.5 | M1/70 | BD, #561114 |
| CD11c | APC | N418 | Biolegend (#117310) |
| CD3 | BV711 | 17A2 | Biolegend, (#100241) |
| CD4 | PE-Cy7 | GK1.5 | Biolegend, # 100422 |
| CD8α | PE | 53-6.7 | Biolegend, # 100708 |
| CD8α | BV510 | 53-6.7 | Biolegend, #100752 |
| CD49b | APC | DX5 | Biolegend, #108910 |
| **LIGHT/HVEM/LTβR/ pathway** |  |  |  |
| LIGHT | Biotin | 3D11 | Home-made (6) |
| HVEM | Biotin | 6C9 | Home-made (8) |
| LTβR | Biotin | 4H8 | Gift from Dr. Carl Ware. |
| **Co-inhibitory molecules** |  |  |  |
| PD-1 | BV421 | 29F.1A12 | Biolegend, #135217 |

ECACC: European Collection of Authenticated Cell Cultures.

### Statistical analysis

Sample normality distribution was analyzed using D’Agostino and Pearson tests. One way ANOVA and a post-analysis based on Tukeýs test was applied to compare the differences of means among groups. The kinetics of tumor growth volume over time for each experimental group was compared using the correction of Holm-Šídák method, which is a statistical approach used to compare multiple pairs of means at different time points. The statistical analysis was performed using GraphPad Prism 8.0 software (GraphPad Software, Inc).

## Results

### Production and purification of two Ig-tagged mouse LIGHT recombinant proteins

LIGHT is a type II transmembrane protein whose C-terminal region contains the trimerization and receptor-binding domain (THD: TNF homology domain). This domain, which can also be shed in a soluble form upon cleavage by a metalloprotease, was fused at the C-terminus of the Fc fragment of mouse IgG_2a_ (hinge, CH2 and CH3 domains) to express an Ig.mLIGHT recombinant protein (Supplementary Figure 2A, 2B, 2C). LIGHT was also expressed as Ig.Foldon-mLIGHT (Supplementary Figure 2D). Foldon is a 27 amino acid-long homotrimeric domain of a T4 phage protein (23). Foldon was inserted N-terminal of mLIGHT with the expectation to stabilize or help folding of mLIGHT as a trimer (Supplementary Figure 2D). Purified Ig.mLIGHT migrated with a size of 52.8 kDa by SDS-PAGE under denaturing and reducing conditions (theoretical Mr 49.3 + 2 x 2.5 kDa for N-linked glycans = 54.3 kDa) and Ig.Foldon-mLIGHT with a size of 59 kDa (theoretical Mr 52.3 + 2 x 2.5 kDa for N-linked glycans = 57.3 kDa). A smaller band of about 25 kDa may correspond to a degradation product (Figure 1).

**Figure 1:**
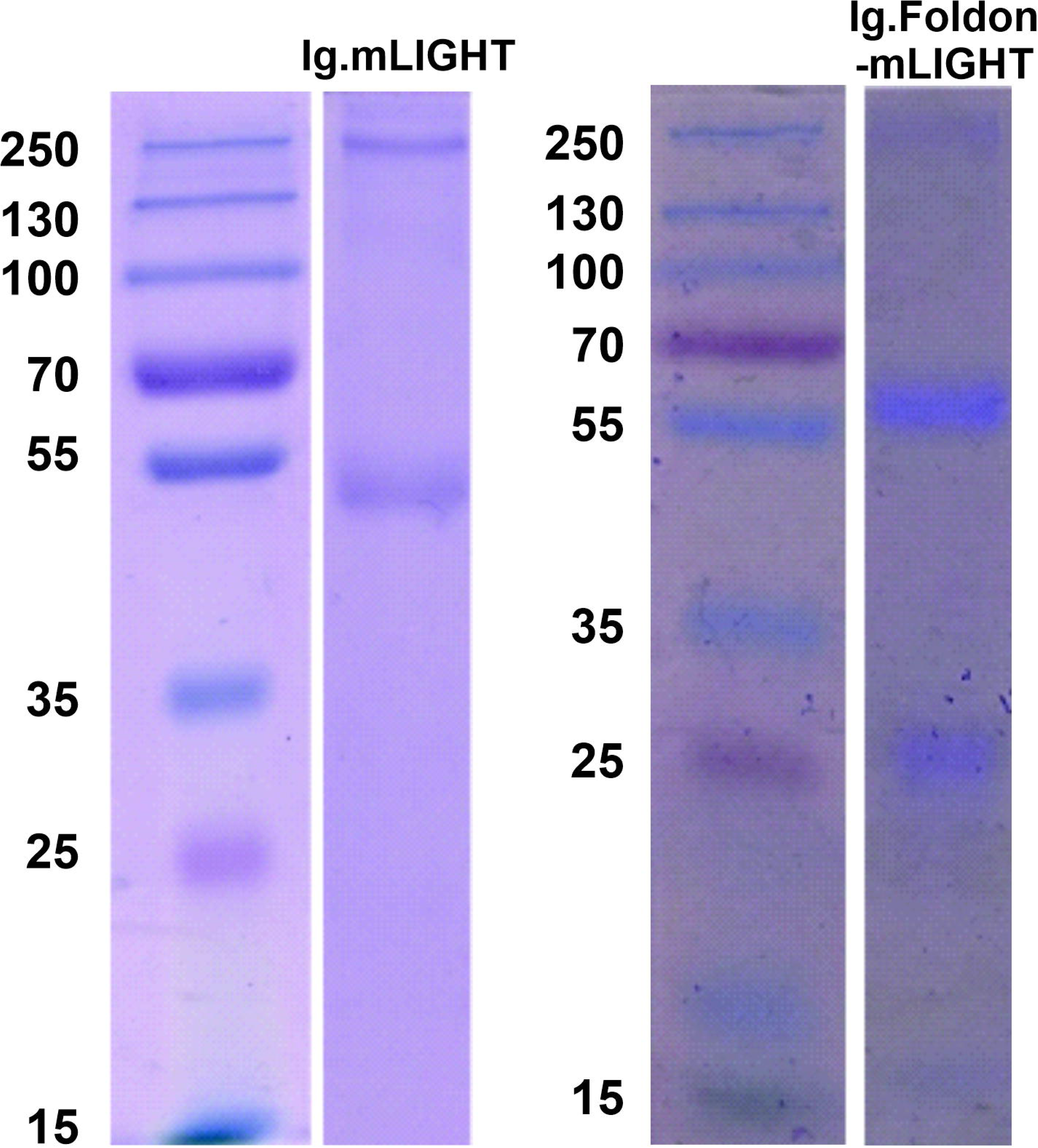
SDS-PAGE electrophoresis of the purified recombinant mouse Ig-LIGHT and Ig.Foldon-LIGHT (ECD) fusion protein. Recombinant soluble mouse LIGHT fusion proteins were purified by Protein G affinity chromatography, as described in material and methods section. Five micrograms of each purified recombinant protein were loaded per lane and run under reduced conditions. Proteins were separated in an 8% resolving SDS-PAGE gel and stained with Coomassie blue. The molecular weight ladder (Thermofisher, page rule plus #11832124) is indicated in the left-hand side of the protein gel.

### The foldon sequence in Ig.Foldon-mLIGHT restored binding to receptors and biological activity

The activity of recombinant LIGHT proteins was assessed using mHVEM:Fas reporter cells. These cells express the extracellular domain of mHVEM fused to the transmembrane and intracellular domain of human Fas, so that addition of active mHVEM ligands kills them through activation of the surrogate Fas death pathway. Recombinant Ig.mLIGHT failed to kill mHVEM:Fas reporter cells, in line with its lack of mHVEM binding (Figure 2A, 2E), but Ig.Foldon-mLIGHT and Flag-hLIGHT killed mHVEM:Fas reporter cells in a dose-dependent manner (Figure 2B, 2F and 2C, 2G). Although soluble mBTLA.Ig could bind mHVEM:Fas in the flow cytometry assay, it could not kill these reporter cells, neither when used as a soluble protein (Figure 2D, 2H) nor when cross-linked via the Fc portion (by a biotinylated polyclonal antibody captured by plate-bound streptavidin, data not shown). Binding of a ligand is therefore not always sufficient to activate mHVEM:Fas. The number and/or the geometry of the receptors in the ligand-induced clusters probably dictate signaling outcome.

**Figure 2:**
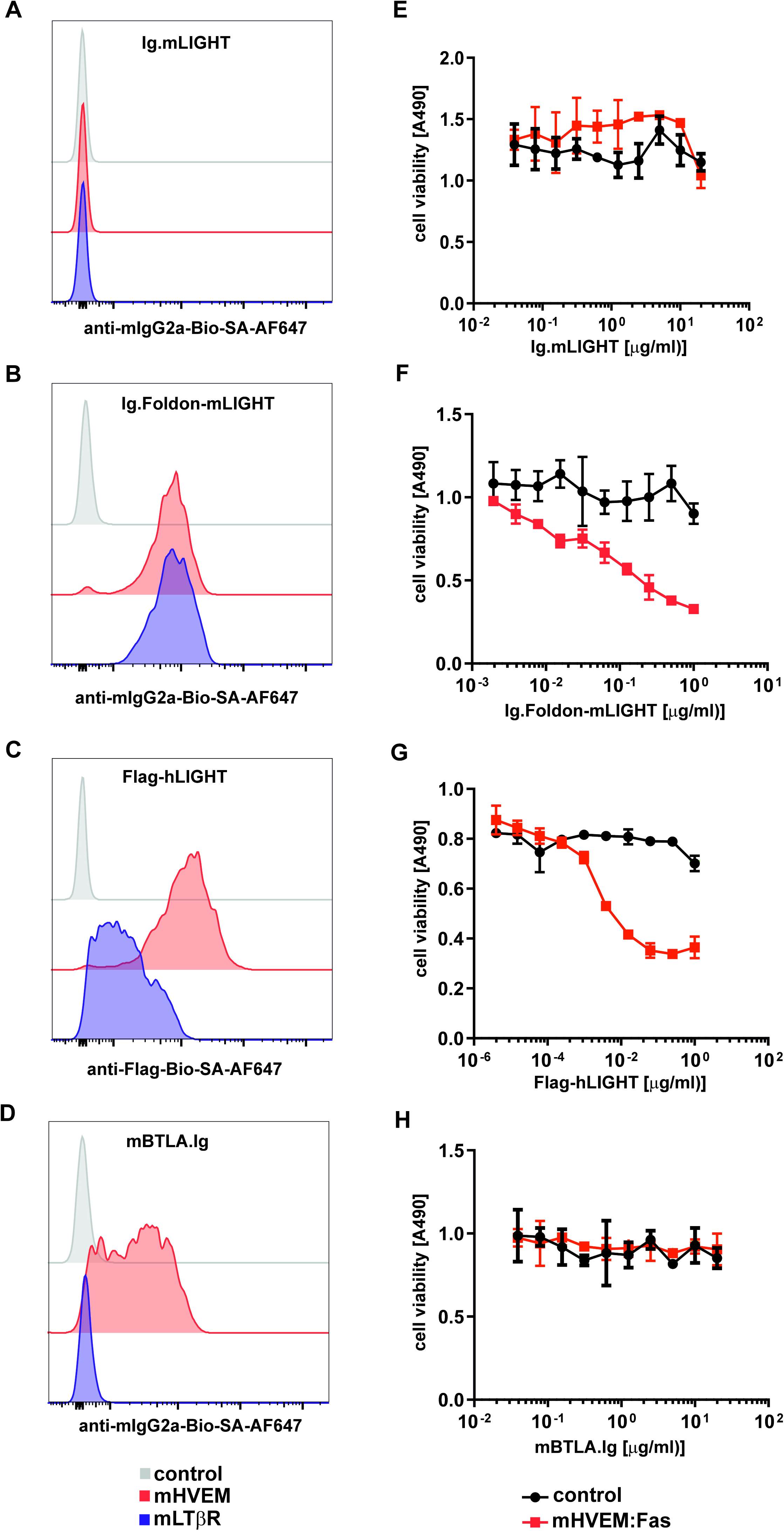
Ig.Foldon-mLIGHT, but not Ig.mLIGHT, binds mHVEM and mLTβR and activates mHVEM:Fas reporter cells. mHVEM:Fas (red lines), mLTβR:Fas (blue lines) and control parental Jurkat cells (grey lines) were stained with recombinant Ig.mLIGHT (**A**), Ig.Foldon-mLIGHT (**B**), Flag-hLIGHT (**C**) or mBTLA.Ig **(D)**. The binding of the fusion protein to the cells was revealed by the addition of a biotinylated anti-mIgG2a **(A, B, D)** or biotinylated anti-Flag antibody **(C**), followed by streptavidin-Alexa Fluor 647. This experiment was performed twice. mHVEM:Fas reporter cells (red squared) and parental Jurkat JOM2 cells (black circles) were exposed overnight to serial dilutions of recombinant Ig-mLIGHT (**E**), Ig.Foldon-mLIGHT (**F**), Flag-hLIGHT (**G**) or mBTLA.Ig (**H**). Cell viability was then measured with MTS/PMS colorimetric test. A representative experiment of two performed is shown.

The specificity of the assay was further demonstrated by preincubation of Ig.Foldon-mLIGHT with an anti-mouse LIGHT blocking antibody (clone 3D11) (6). This function-blocking antibody abrogated cell death induced by Ig.Foldon-mLIGHT and restored reporter cell viability in a dose-dependent manner (Figure 3).

**Figure 3.**
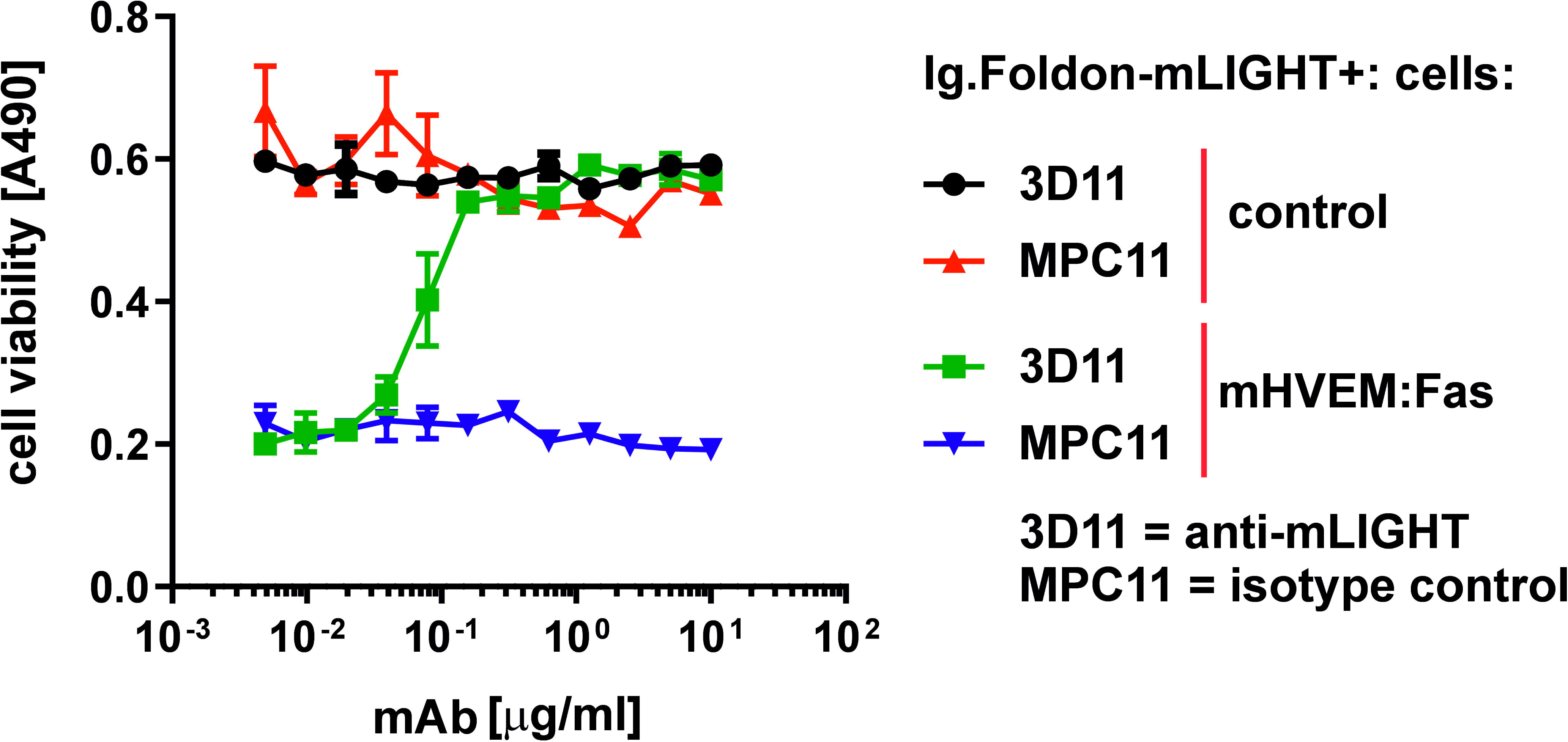
Killing of mHVEM:Fas reporter cells by Ig.Foldon-mLIGHT is blocked by an anti-mLIGHT antibody. mHVEM:Fas reporter cells, or parental Jurakt JOM2 cells used as control, were incubated with a fixed, lethal concentration of 1 µg/ml of Ig.Foldon-mLIGHT preincubated with titrated amounts of an anti-mLIGHT monoclonal antibody (clone 3D11, mouse IgG2b) or of a control antibody (clone MPC11, IgG2b). After an overnight incubation, cell viability was measured with MTS/PMS colorimetric test. A representative experiment of two performed in shown.

In summary, Ig.mLIGHT neither bound its receptors nor activated mHVEM:Fas reporter cells, indicating that mLIGHT may not form stable or properly folded trimers under these conditions. In contrast, Ig.Foldon-mLIGHT bound to mHVEM and mLTβR and was active on reporter cells, possibly because the trimeric foldon helped the folding or arrangement of mLIGHT in an active conformation.

### Superior therapeutic potential of soluble Ig.Foldon-mLIGHT recombinant protein over membrane-bound LIGHT when transduced into B16.F10 melanoma cells

We first isolated a clone (named pIB-176-1F8-5) of B16.F10 melanoma cells transduced with membrane-bound LIGHT by flow cytometry screening using a biotinylated mouse anti-mouse LIGHT monoclonal antibody (clone 3D11) (6) (Figure 4A). Another high expresser clone (named pIB-459b-1B12) secreting soluble Ig.Foldon-mLIGHT was also isolated after transduction of B16.F10 melanoma cells and subsequent cloning by limiting dilution and screening of individual clones by sandwich ELISA. The amount secreted by the selected clone produced 219 ± 9 ng/ml in a period 24 hour after seeding 1×10^6^ cells/ml.

**Figure 4:**
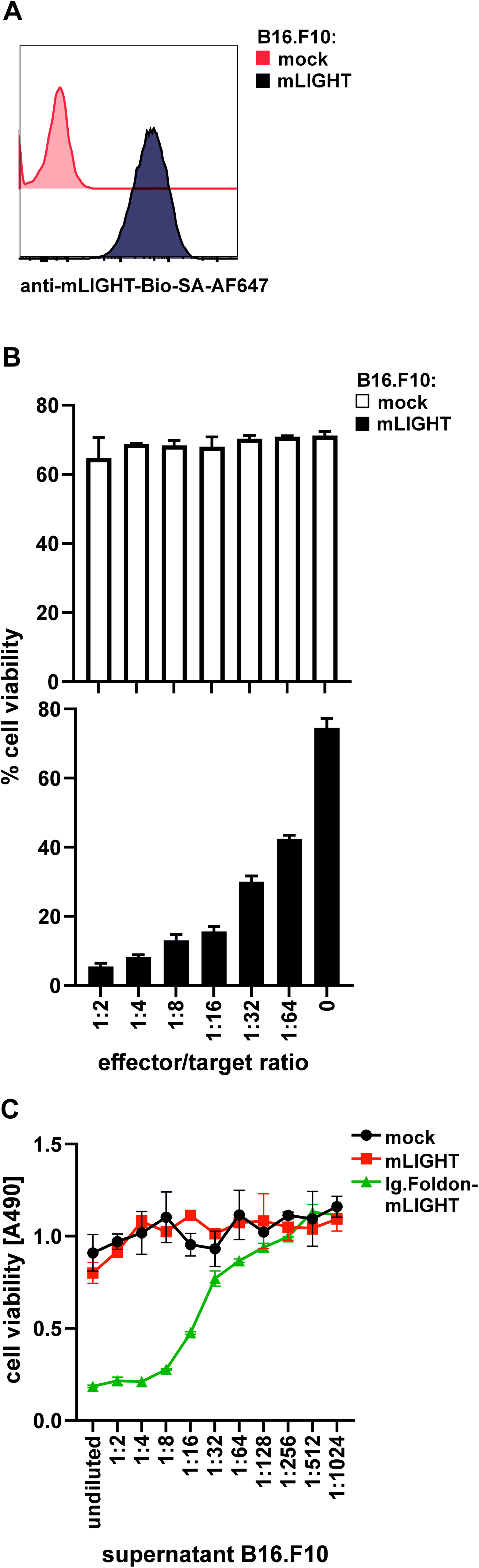
Membrane bound mLIGHT can activate mHVEM:Fas reporter cells. **(A)** Cell surface expression of full-length mLIGHT expressed in a clone of B16.F10 cells (black line) was revealed by flow cytometry after staining with biotinylated anti-mLIGHT antibody (clone 3D11) followed by streptavidin Alexa Fluor 647. Mock control B16.F10 cells (red line) were used as control. **(B)** mHVEM:Fas reporter cells (target) were co-cultured overnight with untransduced B16F10 cells (upper graph) or with a clone of B16F10 cells expressing membrane mLIGHT (effector) at the indicated effector/target ratio (lower graph). Cells viability of non-adherent target cells was measured by flow cytometry. Bars represent mean ± SD of triplicates. One representative experiment of two performed is depicted. **(C)** mHVEM:Fas reporter cells (target) were co-cultured overnight with supernatants from mock B16.F10 cells, membrane mLIGHT B16-F10 cells, or Ig.Foldon-mLIGHT expressing B16.F10 effector cells (1×10^6^ cells/ml grown during 24h). The viability of target cells was measured using the MTS/PMS assay. Symbols represent mean ± SD of duplicate. The experiment was performed once.

Membrane-bound LIGHT expressed on B16.F10 melanoma was functionally active because these cells killed co-cultured mHVEM:Fas reporter cells in an effector to target ratio-dependent manner (Figure 4B). However, conditioned cell supernatants of these cells did not kill reporter cells, indicating that membrane mLIGHT B16.F10 cells do not release soluble mLIGHT, or at least not in an active form. In contrast, conditioned supernatants of B16.F10 cells expressing Ig.Foldon-mLIGHT readily killed reporter cells (Figure 4C).

We next explored whether expression of membrane-bound LIGHT or soluble Ig.Foldon-mLIGHT by tumor cells could inflect tumor growth. Mock control or LIGHT-expressing B16.F10 melanoma cells were implanted into syngeneic mouse recipients and tumor volumes were monitored over time until day 16 post-implantation. There was no difference in tumor growth between control and membrane LIGHT-expressing melanoma cells, even though we hypothesized that biologically active membrane LIGHT on tumor cells could stimulate immune and stromal cells in the microenvironment around the tumor (Figure 5A, B, D). In contrast to membrane LIGHT, tumor growth of melanoma cells expressing Ig.Foldon-mLIGHT was significantly attenuated at multiple time points post-implantation (days 10, 12, 14, and 16) (Figure 5C, D).

**Figure 5:**
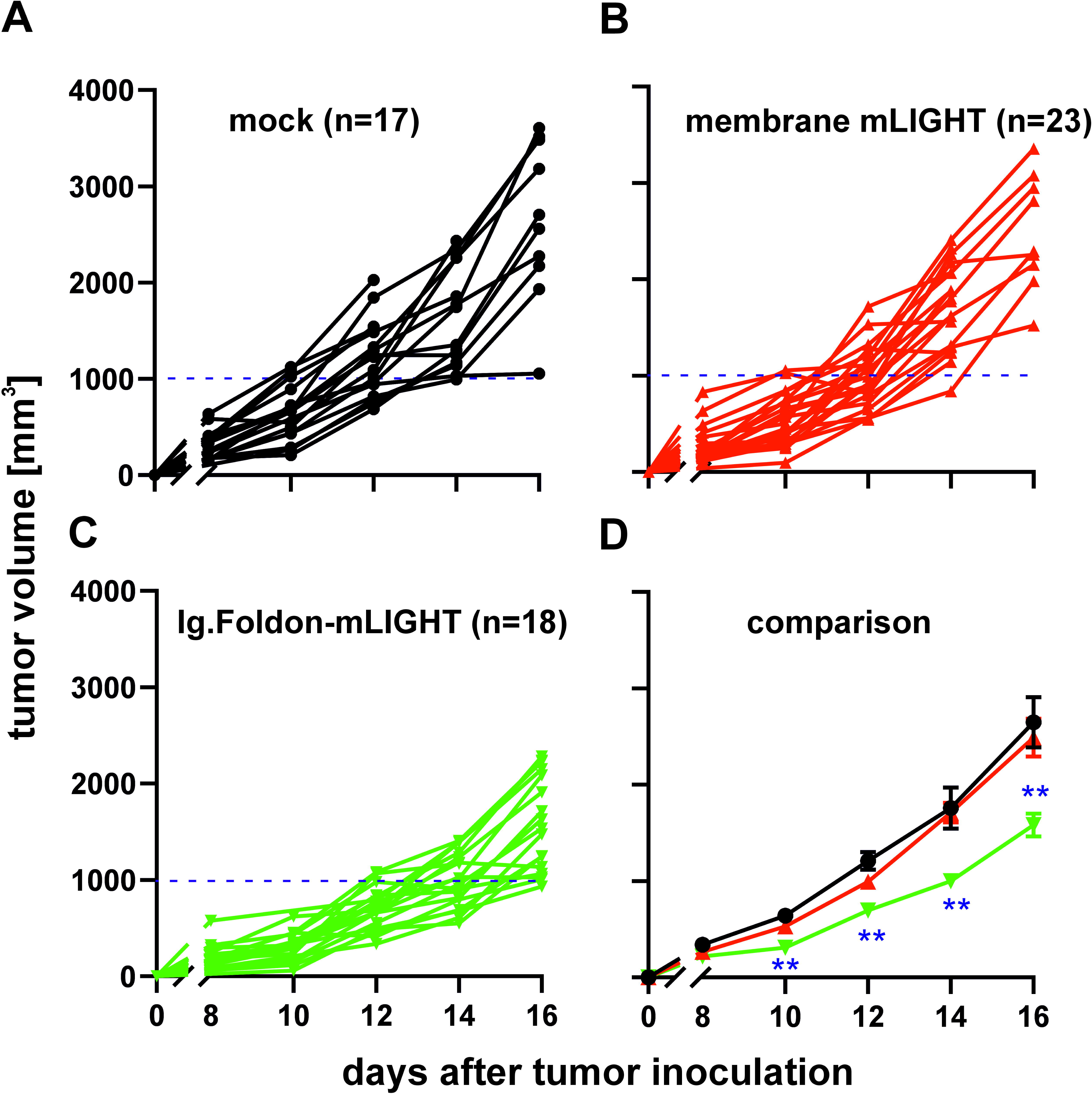
Delayed tumor growth in recipient mice implanted with Ig.Foldon-mLIGHT B16.F10 tumor cells. C57BL/6 mice were injected s.c. with syngeneic mock B16.F10 melanoma cells (n = 17) (**A**), B16.F10 expressing membrane mLIGHT (n = 23) (**B**) or B16.F10 cells secreting Ig.Foldon-mLIGHT (n =18) (**C**). Tumor volume was recorded every other day from day 8 (visible tumors) to day 16. The average tumor volume over time for all three groups ± SEM are compared by multiple testing for each time point with Holm-Sidak correction. * p<0.01, ** p<0.001, *** p<0.001 (**D)**. These plots are a summary of several experiments performed.

Overall, these observations suggest a model in which soluble recombinant LIGHT would act in a paracrine fashion on HVEM expressed on innate and adaptive immune cells- and/or on LTβR-expressing stromal, endothelial and myeloid cells at a distance from the tumor site to trigger a local inflammatory response capable of interfering with tumor growth.

### Delayed tumor growth was associated with increased immune cell infiltration into the tumor microenvironment of Ig.Foldon-mLIGHT B16.F10 tumors

B16.F10 transplantable melanoma is an aggressive tumor model that progresses very quickly over a couple of weeks. Single genetic modifications to express cytokines like Flt3L, GM-CSF, or IP-10 usually do not show any significant anti-tumor activity by themselves unless combined with immune checkpoint blockade or preimmunization strategies with the genetically modified irradiated metabolic active tumor cells before being exposed to live tumor cells (28), (29), (30). This is because B16 melanoma is a poorly immunogenic cold tumor with low expression of MHC class I and with a low rate of tumor immune cell infiltrates. To deepen into the mechanism behind the finding of delayed tumor growth in mice receiving B16.F10 cells expressing Ig.Foldon-mLIGHT, tumor-infiltrating immune cells were counted and phenotypically characterized. There were significantly higher absolute numbers of tumor-infiltrating CD45.2 cells in the Ig.Foldon-mLIGHT group than in the mock control or membrane LIGHT groups (*, p<0.05 and **, p<0.005, respectively) (Figure 6A).

**Figure 6.**
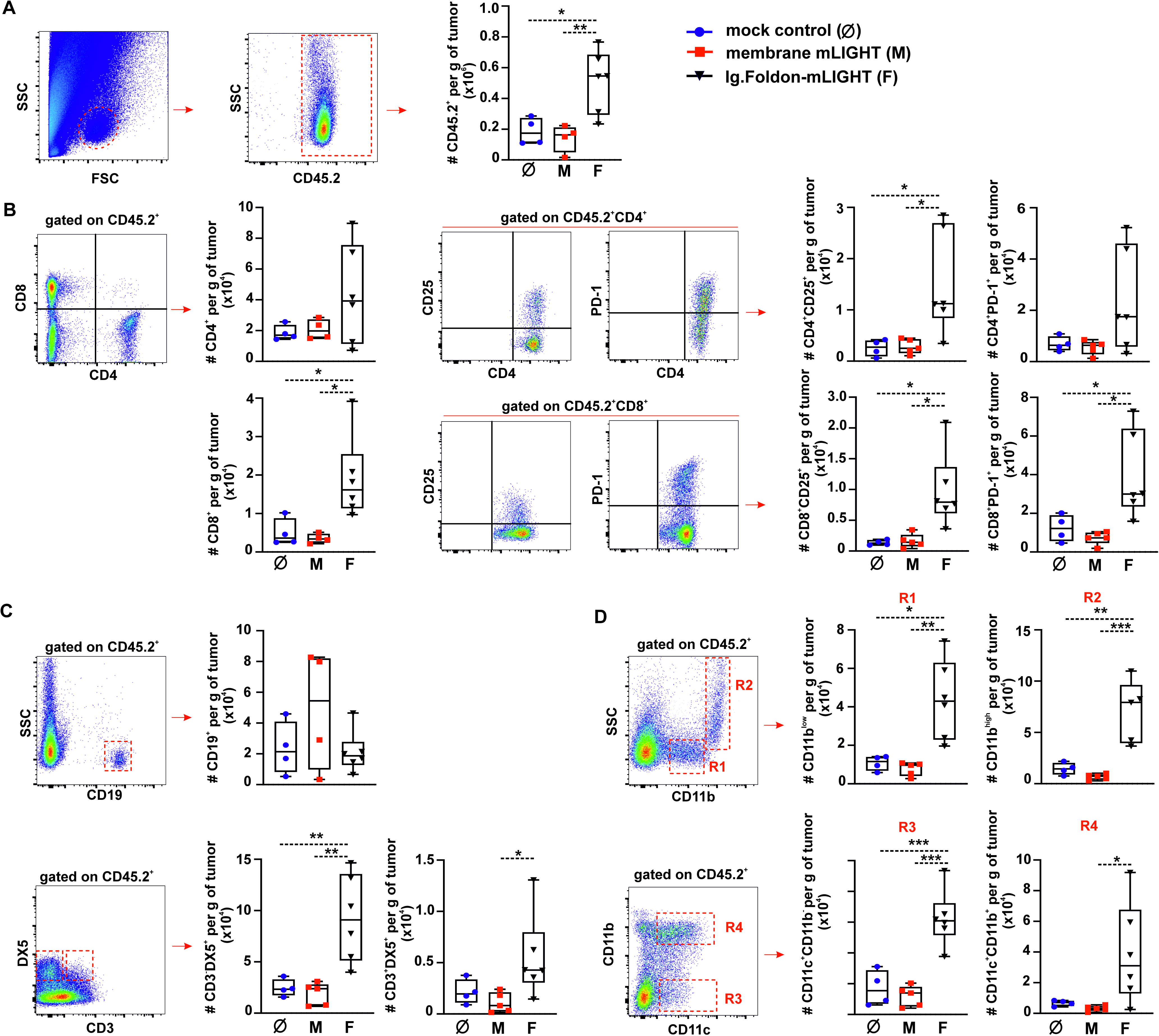
Delayed tumor growth in syngeneic mice implanted with B16.F10 melanoma cells secreting Ig-Foldon-mLIGHT is associated with significant cytotoxic cells and dendritic cells recruitment into the tumor. Tumors of mock control (blue circle, Ø) (n=5), membrane mLIGHT- (red squares, M) (n = 5) or Ig.Foldon-mLIGHT-expressing B16.F10 melanoma cells (black triangles, F) (n = 6) were resected form syngeneic C57BL/6 mice and used to isolate infiltrated immune cells for phenotypic characterization by flow cytometry. (A) FSC/SSC dot plots were used to gate the population of leukocytes. CD45.2^+^ hematopoietic cells were distinguished from CD45.2^−^ non-hematopoietic stromal cells and enumerated. (B) CD4 and CD8 T cells and their subsets of PD-1^+^ and CD25^+^ cells were identified and counted (as number of cells per gram of tumor) by flow cytometry. (C) Same as panel B but for CD19^+^ B cells, CD3^−^/DX5^+^ NK cells and CD3^+^/DX5^+^ NKT cells. (D) Same as panel B but for SSC^low^/CD11b^low^ monocyte-like cells (R1), SSC^high^/CD11b^high^ granulocyte-like cells (R2) and CD11c^+^/CD11b^−^ (R3) and CD11c^+^/CD11b^+^ (R4) dendritic cells. One way ANOVA and a post-analysis based on Tukeýs test was used for the comparisons of means. * p<0.05 ** p<0.005, *** p<0.0005.

We then compared CD4 and CD8 T cells co-expressing either CD25 or PD-1, NK cells (CD3^−^/DX5^+^), NKT cells (CD3^+^/DX5^+^) and B cells (CD19 cells) in the different tumor groups (Figure 6B). In general, the absolute number of immune infiltrating cells was higher in the group of Ig.Foldon-mLIGHT B16.F10 tumors than in the others, with the greatest differences seen in comparison with the group of membrane-bound LIGHT tumors that tended to have less infiltrate than mock control tumors (Figure 6B). CD8 T cells, including CD8^+^/CD25 (IL-2R)^+^ and CD8^+^/PD-1^+^ T cells, CD4^+^/CD25^+^ T cells, NK cells and NKT cells were all significantly higher in the Ig.Foldon-mLIGHT group than in the membrane LIGHT group (and than in the mock control group, except for NKT cells whose increase did not reach significance). There were no statistical differences between groups for CD4 T cells, CD4^+^/PD-1^+^ T cells and for B cells (Figure 6B, 6C).

Finally, distinct populations of myeloid cells in tumor infiltrates were analyzed based on side scatter and the expression of CD11b and CD11c to distinguish CD11b^low^ (low scatter, monocyte-like), CD11b^high^ (high scatter, granulocyte-like), CD11c^+^/CD11b^−^ dendritic cells (conventional cDC1) and CD11c^+^/CD11b^+^ (other DC population). All were elevated in the Ig.Foldon-mLIGHT group compared to the membrane LIGHT group (and also compared to the mock control group, except for CD11c^+^/CD11b^+^ whose increase did not reach significance) (Figure 6D).

The significant presence of various immune cell subpopulations, including myeloid cells and dendritic cells, indicates that LIGHT not only enhances cytotoxic cell activity but also may potentiate antigen presentation and overall immune activation within the tumor microenvironment.

## Discussion

This study reports a recombinant mouse LIGHT protein able to bind its receptors (mouse HVEM and LTβR), to display activity *in vitro* in a cell-based reporter bioassay and, when expressed by tumor cells, to enhance anti-tumor activity *in vivo* in a poorly immunogenic cold tumor model by fostering immune cell infiltration.

Biological assays are of crucial importance for the production and functional characterization of cytokine-like molecules such as native proteins and soluble recombinant forms of members of the TNF superfamily. The measurement of the biological activity of soluble recombinant human LIGHT has relied so far on the use of human tumor cell lines expressing either LTβR and/or HVEM (23), (31), or primary T cells expressing HVEM that can proliferate in response to LIGHT (32). The cell-based bioassay developed in this study for mouse and human LIGHT is convenient, easy to quantify and may discriminate between multimerization states and potencies of the ligands.

With the exception of mouse GITRL that was crystalized as a dimer (33), all other members of the TNF superfamily for which a structure is known, including human LIGHT, have a trimeric configuration and three receptor binding sites. Ligand-induced receptor clustering is at the apex of signaling in the TNF family (1). Approaches commonly used to express active TNF family ligands (*e.g.* Flag-tagged or Ig-tagged) were unsuccessful for mouse LIGHT (2), (23) but insertion of a small homotrimeric domain in front of mouse LIGHT rescued receptor binding and bioactivity, probably by helping mLIGHT folding in an active conformation.

Many patients do not respond to immune check-point blockade, meaning that complementary approaches are needed to overcome tumor resistance to this therapy. Targeting the LIGHT/HVEM/LTβR pathway is gaining attention in the immune-oncology field for several reasons (34), (35), (7). Foundational studies on the biological function of LIGHT showed that the constitutive transgenic LIGHT expression under a T cell promoter leads to an autoimmune syndrome characterized by systemic tissue T cell infiltration and ensuing systemic inflammation (36). Based on this observation, along with other reports claiming anti-tumor activity of membrane LIGHT expressed in immunogenic transplantable tumor cells lines (37), (31), (38), (39), (40), (41), we hypothesized that expression of a recombinant LIGHT could condition the tumor microenvironment and increase anti-tumor responses also in cold, non-immunogenic tumors. Data obtained with B16.F10 melanoma tumor cells expressing Ig.Foldon-mLIGHT support this hypothesis and additionally suggest that mLIGHT can act as a single agent to increase anti-tumor immunity. Previous work with transgenic expression of potent cytokines or chemokines such as GM-CSF, Flt3L, or IP-10 in B16 melanoma cells only displayed anti-tumor activity when combined with immune checkpoint blockade or after prophylactic immunization with irradiated cells expressing these cytokines prior to challenge with live tumor cells (28), (30), but never as single agents as observed with LIGHT. The success of Ig.Foldon-mLIGHT but the failure of membrane LIGHT in the B16.F10 melanoma model helps formulating further hypotheses regarding the mode of action of Ig.Foldon-mLIGHT. Immunogenicity generated by the transduction procedure is unlikely to be a decisive factor as membrane mLIGHT and Ig-Foldon-mLIGHT B16.F10 cells were comparable in that respect. Binding of mLIGHT to receptors, in particular to HVEM is insufficient for the observed anti-tumor effect because membrane LIGHT was active on mHVEM:Fas reporter cells but not on tumors. It is however difficult to compare concentrations and activities of soluble Ig.Foldon-mLIGHT versus membrane LIGHT that might be an intrinsically weaker activator of its cognate receptors. It is also tempting to hypothesize that soluble Ig.Foldon-mLIGHT must act at a distance of the tumor to exert activation and recruitment of immune cells, something that membrane LIGHT cannot do. This hypothesis is reinforced by the observation that membrane mLIGHT B16.F10 cells did not release any active soluble mLIGHT when cultured in vitro.

Ongoing cancer immunotherapy faces several challenges such as the need of promoting anti-tumor responses by bolstering and amplification of pre-existing anti-tumor responses against tumor specific antigens or breaking tolerance to tumor-associated antigens. This therapeutic intervention by itself is insufficient to eliminate solid tumors, since tumor-specific T cells are known to be detectable in peripheral blood, but this is not sufficient since anti-tumor effector cells still need to traffic and infiltrate the tumor parenchyma (42), (43). Once within the tumor, T cells must overcome the immunosuppressive conditions created by regulatory T cells (Tregs) and myeloid-derived suppressor cells for a successful outcome of immune checkpoint blockade (44). The inability of effector cells to migrate into the tumor parenchyma in cold tumors may in part be explained by the observation that tumor vasculature differs from that of normal vasculature (45), (46), (47), (48), (49). Despite the presence of inflammatory conditions, the vascular endothelium paradoxically lacks adhesion molecules (such as selectins and integrins), rendering it incapable of supporting the recruitment of immune cells trafficking through the tumor blood vessel network (47). LIGHT appears to contribute favorably to counteract evasion mechanisms of tumors to subvert the anti-tumor response, thereby enhancing the ability of immune cells to infiltrate the tumor and exert anti-tumor effects (48), (50).

The recombinant expression of LIGHT offers clear advantages over other immunostimulatory molecules as it can influence both immune cell infiltrates and stromal cells of the TME. On the one hand, the LT/LIGHT/HVEM (LTα3/HVEM and LIGHT/HVEM) bidirectional interaction can drive myeloid cell activation, licensing of dendritic cells for antigen presentation, NK cell activation and T cell co-stimulation, thus promoting anti-tumor responses (51), (35), (52), (4). The engagement of HVEM by LIGHT co-stimulates T cells in a CD28-independent manner (14), (19). The engagement of LTβR by LIGHT on DCs seems to compensate for the absence of CD40 ligation in the licensing of DC, a signal required for appropriate T cell co-stimulation (20), (21). Moreover, lymphotoxin and LIGHT can engage LTβR on stromal components of TME, such as fibroblasts and endothelial cells promoting the secretion of chemokines and contributing to endothelial cell activation, accompanied by increased expression of adhesion molecules, resulting at the end in the recruitment of effector cells and the subsequent control of tumor growth (35), (53), (48). The recruitment of both populations of cytotoxic cells, innate NK cells and adaptive CD8 T cells, are required for an effective anti-tumor response. Indeed, NK cells alone do not reject tumors in Rag-deficient mice and NK cell depletion in WT mice prevents effective CTL responses (54), (42). This close link between NK cells, T cells, and DCs and effective clinical anti-tumor responses has been detected intratumorally in patients under the cover of immune checkpoint blockade (55), (56). Our findings are of great significance in the field of immune-oncology as LIGHT can promote beneficial responses through the recruitment and infiltration of the tumor parenchyma of the critical immune effector cells (cytotoxic cells and dendritic cells).

Future studies should address whether combining immune checkpoint blockade and soluble Ig.Foldon-mLIGHT in the B16.F10 model are synergistic approaches capable of eradicating melanoma tumors. Another aspect worth assessing in terms of potential translation to human is to unveil whether human Ig.hLIGHT and Ig.Foldon- hLIGHT similarly designed genetic constructs can induce comparable anti-tumor responses, which should be feasible since human LIGHT can bind mouse LTβR and mouse HVEM. Furthermore, LIGHT-transduced T cells could enhance current adoptive CAR (Chimeric Antigen Receptor) T cell therapy by activating stromal cells and endothelial cells to favor the transmigration and invasion of bystander T cells and CAR T cells to achieve better outcomes in terms of tumor control.

## Supporting information

Graphical abstract

## List of abbreviations

LIGHT: homologous to lymphotoxin, exhibits inducible expression and competes with HSV glycoprotein D for binding to herpesvirus entry mediator, a receptor expressed on T lymphocytes
mAb: monoclonal antibody
CD: Cluster of differentiation
HSV: Herpesvirus
TNF: Tumor necrosis factor
TNFR: Tumor necrosis factor receptor
HVEM: Herpesvirus entry mediator
LTβR: Lymphotoxin β receptor
CRD: Cysteine-rich domain
APC: Antigen-presenting cells
CTL: Cytotoxic T lymphocyte
eGFP: enhanced green fluorescent protein
APC: Allophycocyanin
PE: Phycoerythrin
MHC: Major Histocompatibility Complex
Ig-LIGHT: Immunoglobulin IgG2a Fc-tagged mouse LIGHT
Ig.Foldon-mLIGHT: Immunoglobulin IgG2a Fc (Hinge-CH2-CH2).Foldon-tagged soluble mouse LIGHT
NK: Natural killer
NKT: Natural killer T cells
TCR: T cell receptor
PI: Propidium Iodide
FCS: Fetal calf serum
SD: Standard deviation
PBS: Phosphate buffer saline
BTLA: B- and T-lymphocyte attenuator
LTα: Lymphotoxin alpha
PD-1: Programmed cell death – 1
ECD: Extracellular domain
TNFSF: Tumor necrosis factor superfamily
TNFRSF: Tumor necrosis factor receptor superfamily
CRD: Cysteine rich domains
THD: TNF Homology Domain
TME: Tumor microenvironment
TILs: Tumor-infiltrating lymphocytes
TM: Transmembrane
IC: Intracellular
SA: Streptavidin
AF: Alexa fluor
FSC: Forward scatter
SSC: Side scatter

## Funding and Acknowledgements

This work has been supported by the Spanish Ministry of Science and Innovation # PID2022-136386OB-I00, MCIN/ AEI/10.13039/501100011033/ and “ERDF A way of making Europe” to JIRB and MLRG; Department of Education of Castilla and Leon Regional Government (Grant # LE-087P23) to JIRB and MLRG. P.S. was supported by the Swiss National Science Foundation grant 310030-205196.

We thank Miss Mariana Arroyo Sanchez (technician) and Mr. Israel Esgueva Fuentes and Miss Raquel Alonso Carro (Graduates in Biotechnology) for their technical assistance with cell culture and recombinant molecular biology techniques.

## Authors contributions

M.R.G and J.I.R.B designed and coordinated the study, collected and analyzed data, acquired funding, and wrote the original version of the manuscript. P.S. provided with key scientific inputs, valuable reagents, data interpretation and paper writing. G.R. and J.C. contributed to data interpretation and paper revision. All authors read and contributed to the final version of the manuscript.

## Figure legends

**Supplementary figure 1. Flow chart of the experimental approach**

**(A)** Ig.mLIGHT, Ig.Foldon-mLIGHT and Flag.hLIGHT were purified by affinity chromatography on Protein G-Sepharose or anti-Flag agarose from conditioned supernatants of transiently transfected HEK 293T cells. Binding of the recombinant proteins to mHVEM and mLTβR was assessed by flow cytometry on cells expressing mHVEM:Fas or mLTβR:Fas, and their activity *in vitro* was quantified on mHVEM:Fas reporter cells.

**(B)** HEK293T cells were co-transfected with packaging, envelope and transfer plasmids containing the gene of interest (membrane mLIGHT or Ig.Foldon-mLIGHT) to produce retroviral particles. B16.F10 melanoma cells were transduced with retroviral particles, selected and cloned. Membrane LIGHT expression in B16.F10 melanoma cell clones was screened by flow cytometry with a mouse anti-mouse LIGHT (clone 3D11), while secretion of Ig.mLIGHT and Ig.Foldon-mLIGHT was screened in a sandwich ELISA for the Ig portion of the molecules. Mock control B16.F10 melanoma cells or genetically modified B16.F10 cells were then injected subcutaneously in syngeneic C57BL/6 mice. Tumor volumes and the absolute numbers of tumor-infiltrating immune cells were recorded.

**Supplementary figure 2: mLIGHT, IgG2a and their fusion proteins**

**(A)** Primary structure of mLIGHT with intracellular domain (ICD), transmembrane domain (TMD), the stalk region containing a metalloprotease cleavage site, and the C-terminal extracellular TNF homology domain (THD) containing a N-glycosylation site.

**(B)** Schematic representation of mouse IgG2a, with the hinge and heavy chain constant domains 2 and 3 (CH2 and CH3) used for fusion proteins with mLIGHT. (C) Primary structure of Ig.mLIGHT with the signal peptide of immunoglobulin kappa light chain (signal) fused to hinge, CH2 and CH3 domains of mIgG2a.Fc, itself fused to the THD of mLIGHT (upper). Schematic representation of the putative hexameric Ig.mLIGHT. The irregular shape of the mLIGHT portion symbolizes suboptimal folding or arrangement (lower). (D) Same as panel C, but for Ig.Foldon-mLIGHT, with a short homotrimeric domain (foldon) inserted between the mouse IgG2a.Fc and mLIGHT moieties.

**Graphical abstract**

The intrinsic source of native soluble LIGHT and LTα1β2 (produced by neutrophils and activated T cells and NK cells) along with the tumor secreting recombinant Ig.Foldon-LIGHT co-stimulate NK and T cells through HVEM and license DC for antigen presentation to promote anti-tumor T cell responses in the tumor draining lymph nodes. Furthermore, native LIGHT produced by neutrophils, NK and T cells and recombinant LIGHT along with LTα1β2 secreted by recruited B cells (LTi: lymphoid tissue inducer cells) would together activate stromal cells of the tumor microenvironment (TME) through LTβR (LTo: lymphoid tissue organizing cells) to release chemokines driving endothelial activation and the upregulation of adhesion molecules that in turn would facilitate transmigration of immune cells from the blood vessels to the tumor site. It would also condition myeloid cells expressing LTβR to secrete cytokines such as TGF-beta that contribute to blood vessel normalization.

