## Supplementary figures and images for "A bioactive soluble recombinant mouse LIGHT promotes effective tumor immune cell infiltration delaying tumor growth"

### Graphical abstract

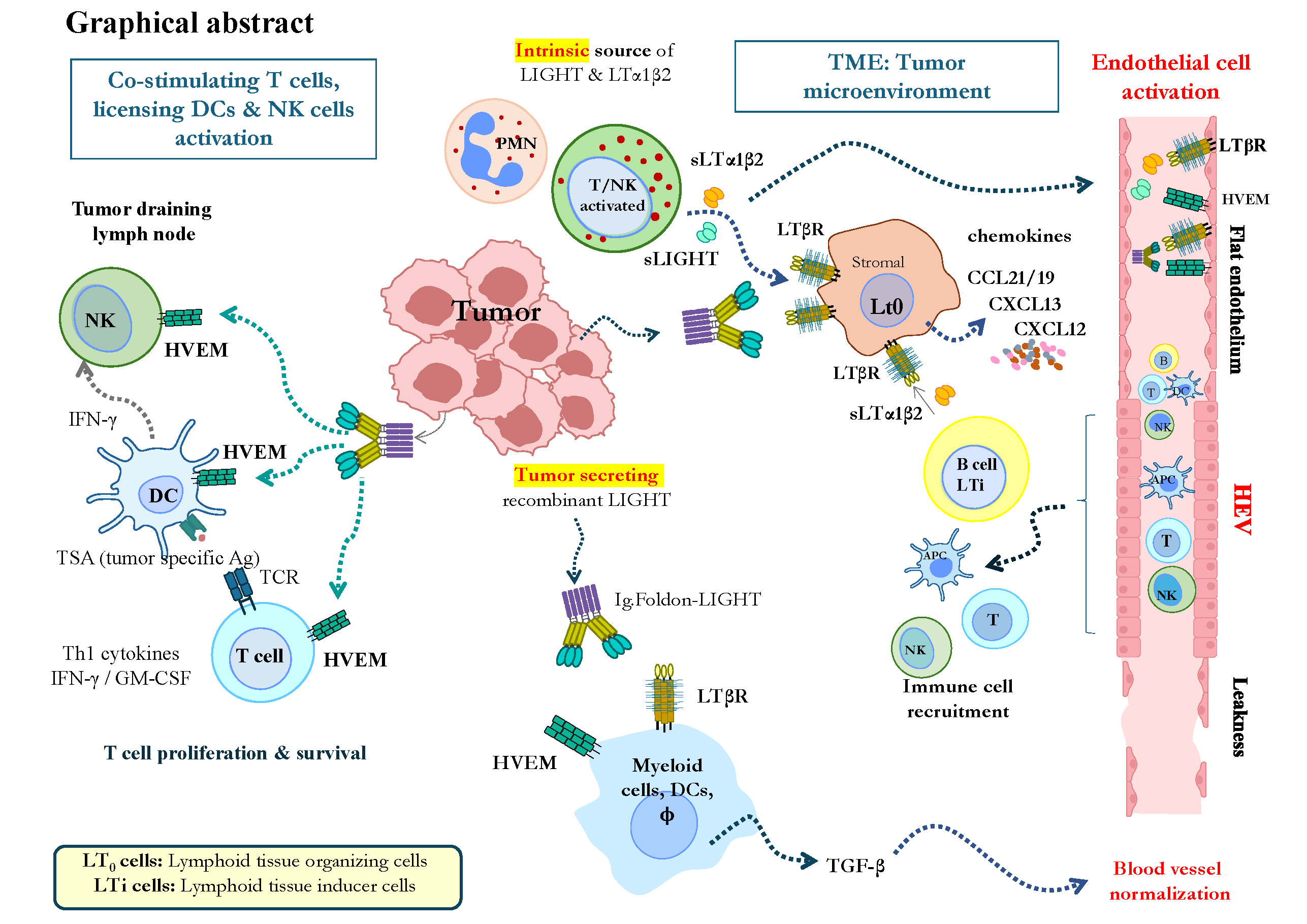

### Supplementary Figure 1

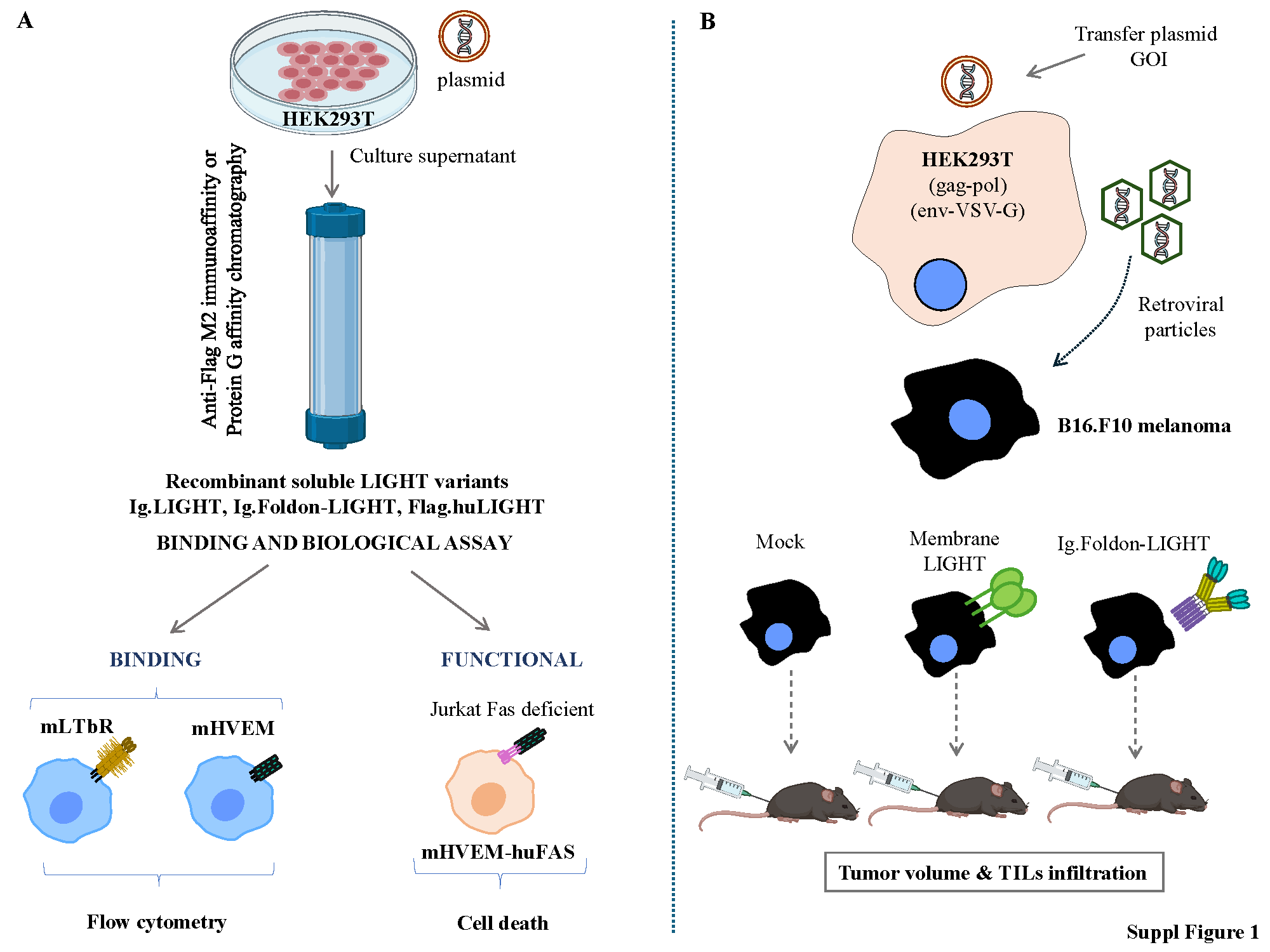

### Supplementary Figure 2

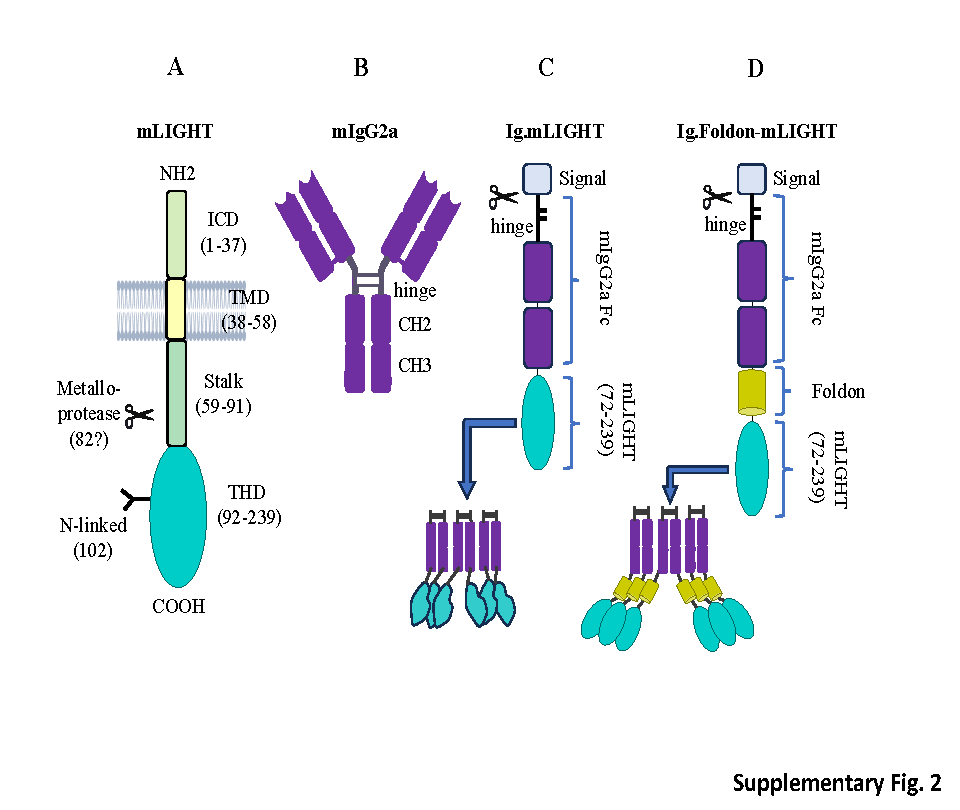
